# Protocol for assessing lysosomal ion channel function in mammalian cells using lysosomal patch-clamp technique

**DOI:** 10.64898/2026.06.01.729450

**Authors:** Yayu Wang, Lily Yeh Jan

**Affiliations:** Department of Physiology, University of California at San Francisco; San Francisco, CA 94158, USA

## Abstract

This protocol describes manual whole-endolysosome patch-clamp recordings from pharmacologically or genetically enlarged endolysosomes in cultured mammalian cells. Steps include vesicle enlargement, fabrication and fire-polishing of high-resistance pipettes, mechanical dissection and isolation of enlarged vesicles, giga-seal formation, and configuration to whole-endolysosome modes. For complete details on the use and execution of this protocol, please refer to Wang et al^1^.

## Before you begin

The following protocol describes the steps required to record endolysosomal chloride currents and to assess lysosomal ion homeostasis in cultured mammalian cells following genetic manipulation of the CLN3 lysosomal chloride channel^2–4^. This protocol is optimized for HEK293T cells and is designed to quantify proton-sensitive Cl^−^ conductance via endolysosomal patch-clamp recording. Nonetheless, the ability to combine high-resolution electrophysiology with live-cell imaging and genetic perturbations enables many additional applications.

This protocol can be used to investigate the impact of disease-associated CLN3 variants on lysosomal channel activity. When combined with complementary assays, such as ion-sensitive fluorescent probes and biochemical measurements, it can be used to examine changes in lysosomal chloride homeostasis and associated effects on lysosomal enzyme activity and protein degradation.

Additionally, this approach has been used to study the regulation of lysosomal chloride levels by lysosomal membrane tension, vesicular trafficking, and autophagy-lysosome pathway dynamics under pharmacological or metabolic stress conditions.

### Innovation

This protocol presents an integrated workflow for directly measuring endolysosomal chloride conductance in intact mammalian cells, representing a significant advance over existing indirect lysosomal ion assays^5–7^. Traditional approaches rely primarily on fluorescence-based reporters or bulk biochemical measurements, which lack the temporal and electrophysiological resolution required to resolve channel-specific activity^8^. Here, we combine endolysosomal patch-clamp electrophysiology with live-cell imaging and genetic perturbation to enable real-time, cell-specific quantification of proton-sensitive Cl^−^ currents.

In addition, customized ionic conditions allow precise control of luminal pH and chloride gradients, facilitating analysis of voltage- and proton-dependent channel gating. The integration of genetic manipulation (e.g., CLN3 variants) with patch-clamp recording further enables rapid structure-function studies and expands applications to lysosomal ion homeostasis and disease mechanisms.

### Preparation of coated coverslips

#### Timing: 2 days (including incubation time)

1. Sterilize microscope glass (12 mm) coverslips by autoclaving before use.
2. Immerse the coverslips in Poly-L-lysine coating solution and incubate for at least 24 h.
3. Remove the coating solution and wash the coverslips thoroughly with double-distilled water (ddH_2_O) five times.
4. Coated coverslips submerged in ddH_2_O can be stored at room temperature for up to one month.
5. Before cell seeding, transfer five coverslips into the 35 mm peri-dish.
6. Allow the coverslips to air-dry completely in a laminar flow hood for at least 15 min prior to use.

**CRITICAL:** Coverslips should be fully dried before application of cells to promote proper attachment.

## Key resources table

| REAGENT or RESOURCE | SOURCE | IDENTIFIER |
| --- | --- | --- |
| Chemicals, peptides, and recombinant proteins |  |  |
| Sodium tetraborate decahydrate | Sigma Aldrich | Cat# S9640 |
| Boric acid | Sigma Aldrich | Cat# B0394 |
| Poly-L-lysine hydrobromide | Sigma Aldrich | Cat# P1399 |
| Vacuolin-1 | Sigma Aldrich | Cat# 673000 |
| YM-201636 | Cayman Chemical | Cat# 13576 |
| Wortmannin | Cayman Chemical | Cat# 10010591 |
| Latrunculin B | Cayman Chemical | Cat# 10010631 |
| DMEM | Thermo Fisher Scientific | Cat# 10566016 |
| Fetal Bovine Serum | Thermo Fisher Scientific | Cat# A5670701 |
| Penicillin-Streptomycin | Thermo Fisher Scientific | Cat# 15140122 |
| JetOPTIMUS DNA transfection reagent | Polyplus | Cat# 101000006 |
| Hydrochloric acid (HCl) | Sigma Aldrich | Cat# 7647-01-0 |
| Methanesulfonic acid (MSA) | Sigma Aldrich | Cat# 471356 |
| N-methyl-D-glucamine (NMDG) | Sigma Aldrich | Cat# M2004 |
| HEPES | Sigma Aldrich | Cat# H3375 |
| MES hydrate | Sigma Aldrich | Cat# M8250 |
| KCl | Sigma Aldrich | Cat# P9541 |
| TrypLE™ express enzyme (1X), no phenol red | Thermo Fisher Scientific | Cat# 12604039 |
| Experimental models: Cell lines |  |  |
| HEK293T | ATCC | Cat# CRL-3216;<br>RRID:CVCL_0063 |
| Recombinant DNA |  |  |
| pcDNA3.1-CLN3-GFP | Thomas Braulke | Addgene Cat# 78110 |
| pmCherry-C3-Rab5 | This paper | N/A |
| pEGFP-N1-Lamp2 | This paper | N/A |
| pSin-EF2-Rab7-puro | This paper | N/A |
| Fiji/ImageJ | National Institutes of Health | https://imagej.net/Fiji |
| Excel | Microsoft Office | https://www.microsoft.com/en-us/ |
| Clampfit | Molecular Devices | https://www.moleculardevices.com/ |
| OriginPro 2018 | Originlab | https://www.originlab.com/ |
| Other |  |  |
| Patch clamp amplifier: Multiclamp 700B | Molecular Devices | N/A |
| Digitizer: Digidata 1550B | Molecular Devices | N/A |
| Multi-micromanipulator systems: MPC-200-ROE | Sutter Instrument | N/A |
| Micropipette puller: Sutter P-1000 | Sutter Instrument | N/A |
| Glass: borosilicate capillaries | Sutter Instrument | Cat# BF150-75-10 |
| Inverted microscope: Olympus IX73 | Olympus | N/A |
| Osmometer: Osmotech | Advanced Instrument | N/A |
| Microforge: Narishige | Sutter Instrument | N/A |

## Materials and equipment

| Vacuolin-1 stock solution (1 mM) |  |  |
| --- | --- | --- |
| Reagent | Final concentration | Amount |
| Vacuolin-1 | 1 mM | 1 mg |
| DMSO | N/A | 1.73 mL |
| Total | N/A | 1.73 mL |

**CRITICAL:** Prepare Vacuolin-1 stock solution in an appropriate solvent according to the manufacturer’s solubility information (e.g., DMSO). Aliquot the solution after preparation to prevent product inactivation caused by repeated freeze-thaw cycles.

**CRITICAL:** Vacuolin-1 stock solutions can be stored at −80 °C for up to 6 months or at −20 °C for up to 1 month (stored under nitrogen).

| YM201636 stock solution (1 mM) |  |  |
| --- | --- | --- |
| Reagent | Final concentration | Amount |
| YM201636 | 1 mM | 1 mg |
| DMSO | N/A | 2.139 mL |
| Total | N/A | 2.139 mL |

**CRITICAL:** YM201636 stock solutions can be stored at −80°C for long-term use or at −20°C for short-term use. Avoid moisture exposure and ensure the solution is fully dissolved before use.

| Wortmannin stock solution (1 mM) |  |  |
| --- | --- | --- |
| Reagent | Final concentration | Amount |
| Wortmannin | 1 mM | 1 mg |
| DMSO | N/A | 2.3343 mL |
| Total | N/A | 2.3343 mL |

**CRITICAL:** Wortmannin is unstable in aqueous solution and sensitive to moisture, light, and temperature. Stock solutions should be prepared in anhydrous DMSO, aliquoted to avoid repeated freeze-thaw cycles, and stored at −80°C for long-term use or −20°C for short-term use. Handle under minimal light exposure and limit air contact. Ensure complete dissolution before use, and prepare fresh working solutions prior to each experiment.

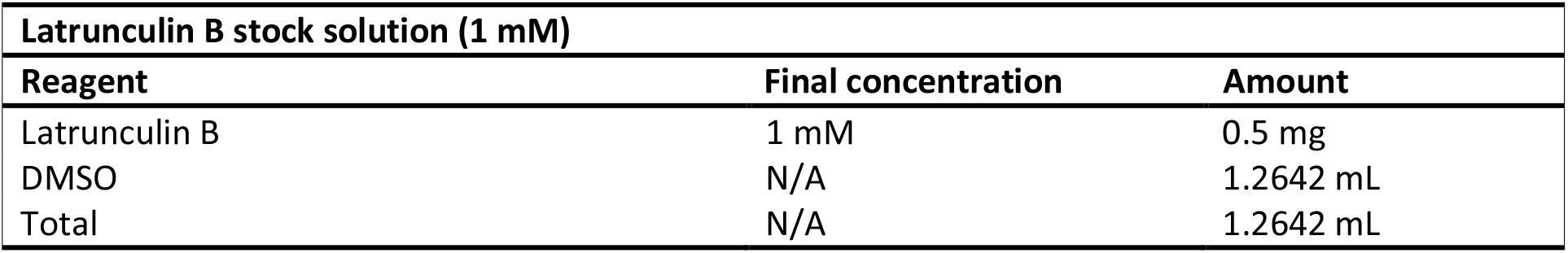

**CRITICAL:** Latrunculin B stock solutions can be stored at −80°C for long-term use or at −20°C for short-term use. Avoid moisture exposure and ensure the solution is fully dissolved before use.

| Poly-L-lysine coating solution (0.1 mg/mL) |  |  |
| --- | --- | --- |
| Reagent | Final concentration | Amount |
| Poly-L-lysine | 0.1 mg/mL | 1 mg |
| Boric acid | 80 mM | included in buffer |
| Borate | 10 mM | included in buffer |
| ddH<sub>2</sub>O | N/A | 10 mL |
| Total | N/A | 10 mL |

**CRITICAL:** The pH of the borate buffer should be carefully adjusted (typically 8.5), as it can significantly affect the efficiency of Poly-L-lysine coating.

**CRITICAL:** The PLL solution is sterilized by filtration (0.22 µm).

### Pipette solution

Pipette solution is used as the intracellular recording solution for electrophysiological experiments. Prepare the solution in advance and store at −20°C until needed. Before use, allow the solution to reach room temperature. Adjust the pH to 4.6 and osmolarity to 300 mOsm.

| Reagent | Final concentration | Amount |
| --- | --- | --- |
| HCl | 150 mM | 625 μL |
| NMDG | 150 mM | 1.46 g |
| HEPES | 10 mM | 0.119 g |
| MES | 10 mM | 0.0976 g |
| ddH<sub>2</sub>O | N/A | 49.375 mL |
| Total | N/A | 50 mL |

**CRITICAL:** Concentrated hydrochloric acid is corrosive. Handle with appropriate personal protective equipment and prepare the solution in a chemical fume hood. Add acid slowly to water while mixing.

**CRITICAL:** The pipette solution is sterilized by filtration (0.22 µm).

### Bath solution

Bath solution is used as the extracellular recording solution during electrophysiological experiments. Prepare the solution in advance and store at 4 °C until needed. Before use, allow the solution to reach room temperature. Adjust the pH to 7.2 and osmolarity to 300 mOsm.

| Reagent | Final concentration | Amount |
| --- | --- | --- |
| Methanesulfonic acid | 149 mM | 4.84 mL |
| HCl | 1 mM | 41.7 μL |
| NMDG | 150 mM | 14.64 g |
| HEPES | 20 mM | 2.38 g |
| ddH<sub>2</sub>O | N/A | 495.11 mL |
| Total | N/A | 500 mL |

**Note:** Dissolve the reagents in 80% of the final volume of ddH_2_O, then bring the solution to the final volume with additional ddH_2_O.

| HEK293T medium |  |  |
| --- | --- | --- |
| Reagent | Final concentration | Amount |
| DMEM high glucose (contained GluMAX) | N/A | 445 mL |
| FBS | 10 % (v/v) | 50 mL |
| Penicillin-streptomycin (100x) | 1 x | 5 mL |
| Total | N/A | 500 mL |

**CRITICAL:** The complete medium should be stored at 4 °C and warmed to 37 °C before use.

## Step-by-step method details

### Preparation of HEK293T cells and transfection for endolysosomal recordings

#### Timing: 2-4 days

1. Maintain cells
  a. Keep HEK293T cells in supplemented DMEM at 37 °C in a humidified incubator with 5% CO_2_.
  b. Passage cells every 2-3 days to maintain healthy growth.

**NOTE:** HEK293T cells should be passaged before reaching 80-90% confluency to maintain healthy growth.

2. Cell detachment
  a. Remove culture medium from a T25 flask.
  b. Add 1.0 mL of TrypLE™ express enzyme. Incubate at 37 °C, 5% CO_2_ for 2 min.
  c. Check detachment under a microscope. If cells remain attached, incubate for an additional 1 min.

**NOTE:** Avoid over-digestion, as prolonged exposure to TrypLE™ may reduce cell viability.

3. Quench trypsin
  a. Add 4.0 mL of supplemented DMEM to dilute trypsin.
  b. Pipette the suspension up and down gently to ensure a single-cell suspension.

**NOTE:** Gentle pipetting minimizes cell damage and improves uniform seeding.

4. Cell seeding
  a. Transfer 0.5 mL of cell suspension (8 × 10^5^ cells) to a new 35-mm peri-dish with 1.5 mL of supplemented DMEM.
  b. Incubate at 37 °C, 5% CO_2_ for 1-2 days until about 80% confluent.

**NOTE:** Optimal confluency (70-80%) is critical for achieving consistent transfection efficiency.

5. Transfection (1-2 days before patch-clamp)
  a. Transfect cells with plasmid DNA encoding CLN3 using polyplus following the manufacturer’s instructions. For a 35 mm peri-dish, mix 2.0 μg plasmid DNA (EGFP-CLN3) with 200 μL buffer, then add 4 μL Polyplus reagent, mix gently and incubate for 10 min at room temperature. Add the transfection mix dropwise to cells in 2.0 mL complete DMEM.

**NOTE:** Use a single transfection reagent consistently throughout the study to ensure reproducibility.

**CRITICAL:** Fluorophore-tagged endolysosomal channel proteins enable visualization and identification of endolysosomes, facilitating whole-endolysosome patch-clamp recordings.

b. Incubate cells at 37 °C, 5% CO_2_ for 24 h.
c. After 24 h post-transfection, detach and re-plate as Step 2-3 for vesicle enlargement and recordings. Verify transfection efficiency by EGFP fluorescence before proceeding.

**NOTE:** Re-plating improves cell morphology and accessibility for patch-clamp recordings.

### Enlargement of endolysosomal vesicles

#### Timing: 1-48 h (depends on reagent and cell type)

This step induces enlargement of late endosomes and lysosomes to facilitate imaging, vesicle tracking, and functional analysis of endolysosomal compartments (Figure 1).

**Figure 1.**
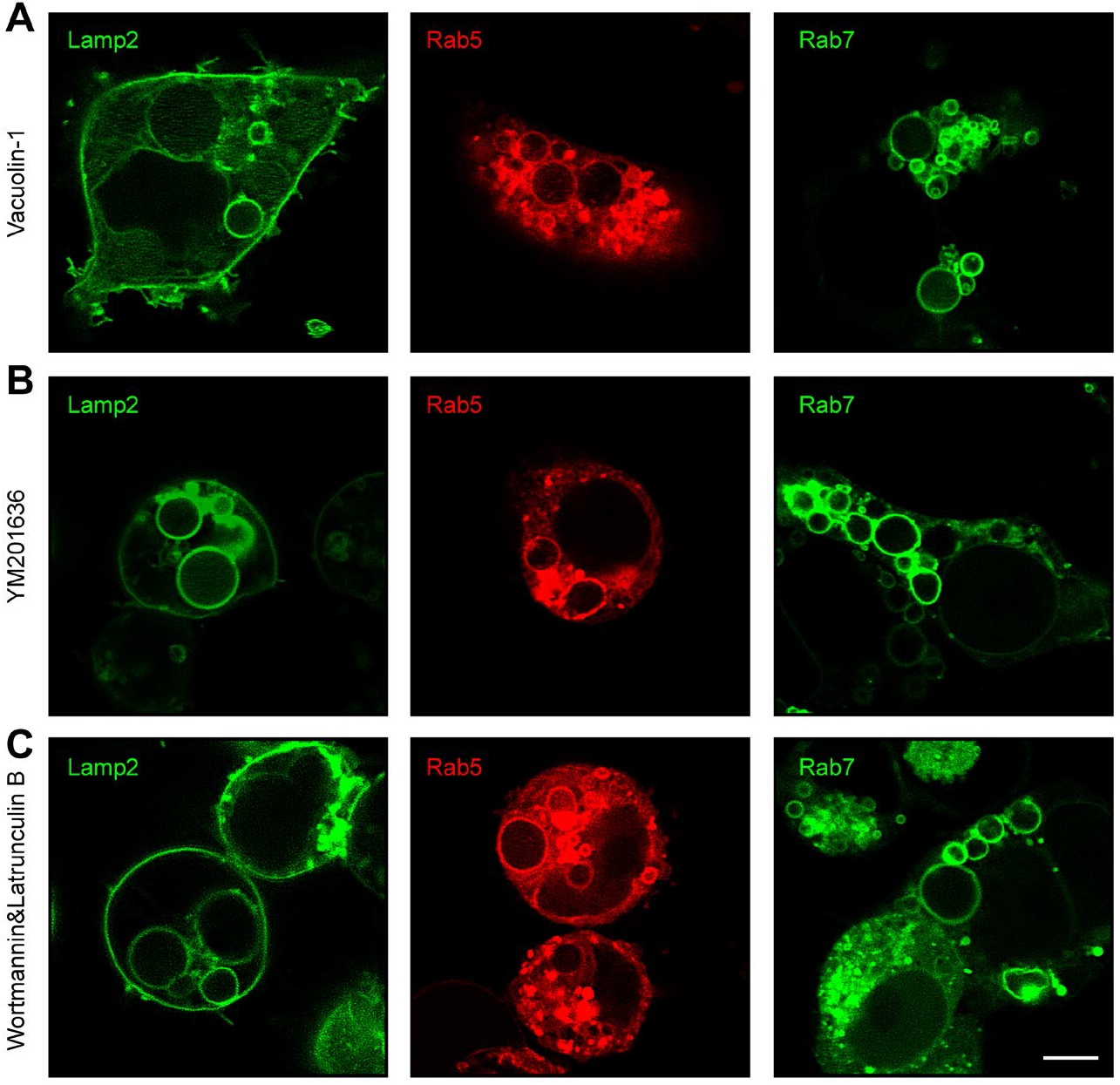
Pharmacological perturbation alters endolysosomal size. Representative confocal images of cells stained for Lamp2 (green), Rab5 (red), and Rab7 (green) under the indicated treatments: (A) Vacuolin-1, (B) YM201636, and (C) Wortmannin in combination with Latrunculin B. Scale bar, 5µm.

6. Treat cells with Vacuolin-1, YM-201636 or Wortmannin/Latrunculin B to induce endolysosomal enlargement.
  a. For Vacuolin-1 treatment^9^:
    i. Add Vacuolin-1 to a final concentration of 1 µM and incubate overnight.
    ii. Alternatively, use 1-5 µM for 1 h in certain cell types.

**NOTE:** Vacuolin-1 induces relatively non-specific enlargement of endolysosomal compartments. The optimal concentration and incubation time may vary depending on the cell type.

b. For PIKfyve inhibition using the YM-201636 compound^10^:
  i. In HEK293T cells, treat with 0.4 µM overnight or 0.8 µM for 2 h.
  ii. In macrophages, treat with 0.4 µM for 1-3 h.

**NOTE:** YM-201636 specifically inhibits PIKfyve and preferentially enlarges late endosomes and lysosomes. Treatment conditions should be optimized to balance enlargement efficiency and cellular health.

c. For Wortmannin + Latrunculin B treatment:
  i. In HEK293 or COS-1 cells, treat with 200 nM Wortmannin + 10 nM Latrunculin B overnight.
  ii. Alternatively, use 100-500 nM Wortmannin + 1-2 µM Latrunculin B for 30-60 min in certain cell types.

7. Confirm vesicle enlargement by live-cell imaging.
  a. Use LAMP or Rab markers to visualize and verify enlargement of late endosomes/lysosomes.
  b. Optimize concentration and incubation time for each cell type.
  c. Compare different treatment conditions to rule out drug-induced artifacts.

**NOTE:** The efficiency and extent of vesicle enlargement depend strongly on cell type, reagent concentration, and incubation time.

### Fabrication and fire-polishing of patch pipettes

#### Timing: 1-10 min per pipette (pulling) + 1-3 min polishing

The fabrication of glass pipettes is a critical step for successful endolysosomal patch-clamp recordings, as optimal pipette geometry and resistance are required for giga-seal formation on fragile intracellular organelle membranes.

8. Mount the glass capillary (e.g., BF150-75-10) onto the puller according to the manufacturer’s instructions.
9. Run a RAMP test to determine the appropriate HEAT value required to melt the glass under current experimental conditions.

**CRITICAL:** The HEAT parameter depends on filament condition, room temperature, and glass type. Recalibration is required if the filament ages or experimental conditions change.

10. Program a six-cycle pulling protocol using optimized HEAT, velocity (VEL), and time (TIME) parameters in given Figure 2. Set HEAT based on Step 9.

**Figure 2.**
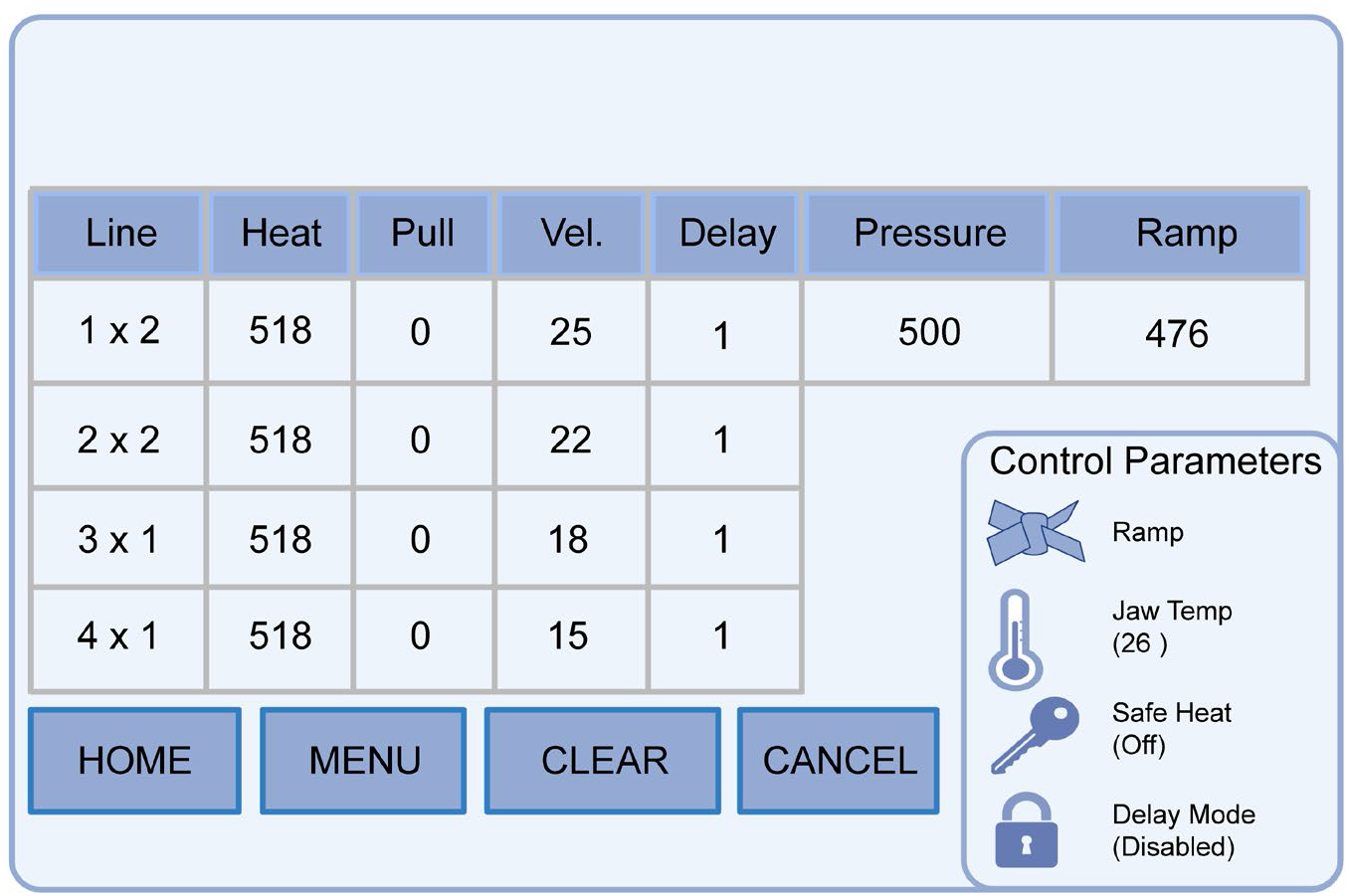
Patch-clamp recording parameters for lysosomal electrophysiology. Representative configuration of acquisition parameters used for lysosomal patch-clamp recordings, including heat, pull, velocity (Vel.), delay, pressure, and ramp settings. Control parameters for ramp, jaw temperature, safe heat, and delay mode are shown on the right panel.

**NOTE:** Pulling parameters must be optimized for each puller and glass type; values are not universally transferable.

**NOTE:** We use a Sutter P-1000 puller with a six-cycle program, which provides high reproducibility for endolysosomal patch pipettes.

11. Initiate the pulling program. Use six-cycle pulling program optimized for endolysosomal pipettes (aim for final pipette resistance 5-8 MΩ).

**CRITICAL:** Capillary separation should occur during the sixth cycle. Earlier separation typically produces overly sharp tips that are unsuitable for giga-seal formation.

12. Remove pipettes from the puller.
13. Inspect pipette tips under a microforge to assess tip diameter, shape, and symmetry. Discard obviously defective tips. Fire-polish remaining pipettes to obtain blunt tip geometry with inner path narrow and tip opening <0.9 µm; target final tip inner diameter <0.3 µm after polishing. Desired tip opening: 0.5-0.9 μm.

**NOTE:** Pipette resistance (typically 5-8 MΩ) is primarily determined by tip geometry, glass properties, and internal solution composition.

**CRITICAL:** Proper pipette geometry is essential for achieving stable giga-seal formation on endolysosomal membranes. An excessively wide (>1 μm) or thick pipette tip can be corrected by increasing the HEAT parameter (sixth cycle) or the pulling velocity (VEL). In contrast, overly narrow or thin tips can be adjusted by reducing the HEAT or VEL parameters accordingly.

**CRITICAL:** Fine adjustment of HEAT and VEL (especially in the sixth cycle) is required for each pulling session.

14. Discard unsuitable pipettes (too sharp, too wide, too thin, asymmetric, or contaminated with debris).

**CRITICAL:** Approximately 50% of pipettes may fail quality control and should not be used for recordings.

**NOTE:** Suboptimal pipettes may be repurposed for dissection or other applications.

15. Immediately fire-polish suitable pipettes using a microforge.

**CRITICAL:** Fire-polishing is essential to produce a smooth, slightly blunt tip, which improves seal formation and minimizes membrane damage.

**CRITICAL:** Use thick-walled glass; produce many pipettes (≥10-20) because 50% of those pipettes fail QC. Use polished pipettes within 6 h.

### Dissection and release of enlarged endolysosomes

#### Timing: 1-10 min per vesicle

16. Transfer coverslip with enlarged cells to the recording chamber and add 1 mL bath solution. Keep cells stable and perform dissections within 1 h of transfer.

**CRITICAL:** Perform dissections within 1 h after transfer; prolonged incubation leads to vesicle shrinkage, increased adhesion, and cell detachment.

17. Fill an isolation pipette with luminal solution and mount it on the headstage. Approach the cell at about 30° using the micromanipulator and select vesicles located near the cell edge and of appropriate size.

**CRITICAL:** Vesicles near the cell edge are easier to isolate; their unsuitable positioning significantly reduces success rate.

**NOTE:** Vesicles should be <50% of the cell diameter; overly large vesicles are unstable, whereas smaller vesicles are difficult to patch.

18. Bring the pipette into contact with the plasma membrane near the target vesicle. Press gently and rapidly withdraw to create a membrane opening.

**CRITICAL:** Use rapid but controlled movement; slow or imprecise withdrawal reduces successful membrane incision.

19. Use the same pipette to push from the opposite side of the cell and squeeze the enlarged endolysosome out through the incision. Move the isolated vesicle away from the cell to avoid reattachment or contamination.

**CRITICAL:** Apply minimal and controlled force; excessive pressure may rupture the vesicle.

**NOTE:** Isolated vesicles may transiently remain attached or drift back toward the cell.

20. Allow the vesicle to stabilize and confirm morphology (round shape, intact, thin and semi-transparent membrane) before proceeding to patch-clamp recording.

**CRITICAL:** Proceed only with intact vesicles; damaged membranes will not form stable giga-seals.

**NOTE:** Vesicles should appear round with a thin, semi-transparent membrane.

### Giga-seal formation

#### Timing: 1-3 min for giga-seal; break-in <1 min

21. Use a freshly polished recording pipette and fill it with the appropriate luminal solution.

**CRITICAL:** Use freshly polished pipettes (<6 h); aged or contaminated tips significantly reduce seal success.

22. Mount the pipette onto the headstage of the amplifier.
23. Apply positive pressure (0.05-0.1 mL) using a 1 mL syringe and maintain constant pressure.

**CRITICAL:** Positive pressure prevents clogging and improves seal formation. Excessive pressure (>0.1 mL) may hinder subsequent break-in for whole-endolysosome configuration.

24. Set the micromanipulator MODE to 0. Lower the pipette into the bath and position it in the center of the visual field. Confirm visible outflow from the pipette tip.

**NOTE:** Visible fluid flow indicates proper pressure and helps keep the tip clean before contacting the vesicle.

25. Apply repetitive test pulses (+5 mV, 50-100 ms) to monitor pipette resistance and seal formation.

**NOTE:** A decrease in current amplitude indicates increasing seal resistance during giga-seal formation.

26. Set MODE to 6 and rapidly move the pipette to the top of the target vesicle (<30 s).

**CRITICAL:** Approach quickly but smoothly; prolonged exposure increases contamination risk and reduces seal probability.

27. Advance the pipette until vesicle movement is observed due to fluid flow.

**NOTE:** Slight vesicle displacement indicates correct positioning and appropriate pressure.

28. Adjust the pipette offset potential to 0 mV.
29. Release the positive pressure immediately. The vesicle membrane should be drawn to the pipette tip and form a giga-seal (1-20 GΩ) within 1 s.

**CRITICAL:** Timing of pressure release is essential; delayed or gradual release reduces giga-seal formation efficiency.

**CRITICAL:** Only intact, stable vesicles will form high-resistance seals.

### Recording configurations after giga-seal formation

After formation of a stable giga-seal, proceed with the desired recording configuration as outlined below.

#### Timing: 1-5 min, depending on the selected recording configuration (Options A-D) (Figure 3)

A. **Endolysosome-attached mode (single-channel recording)** Maintain the giga-seal without membrane rupture and proceed directly to single-channel recording using step protocols.

**Figure 3.**
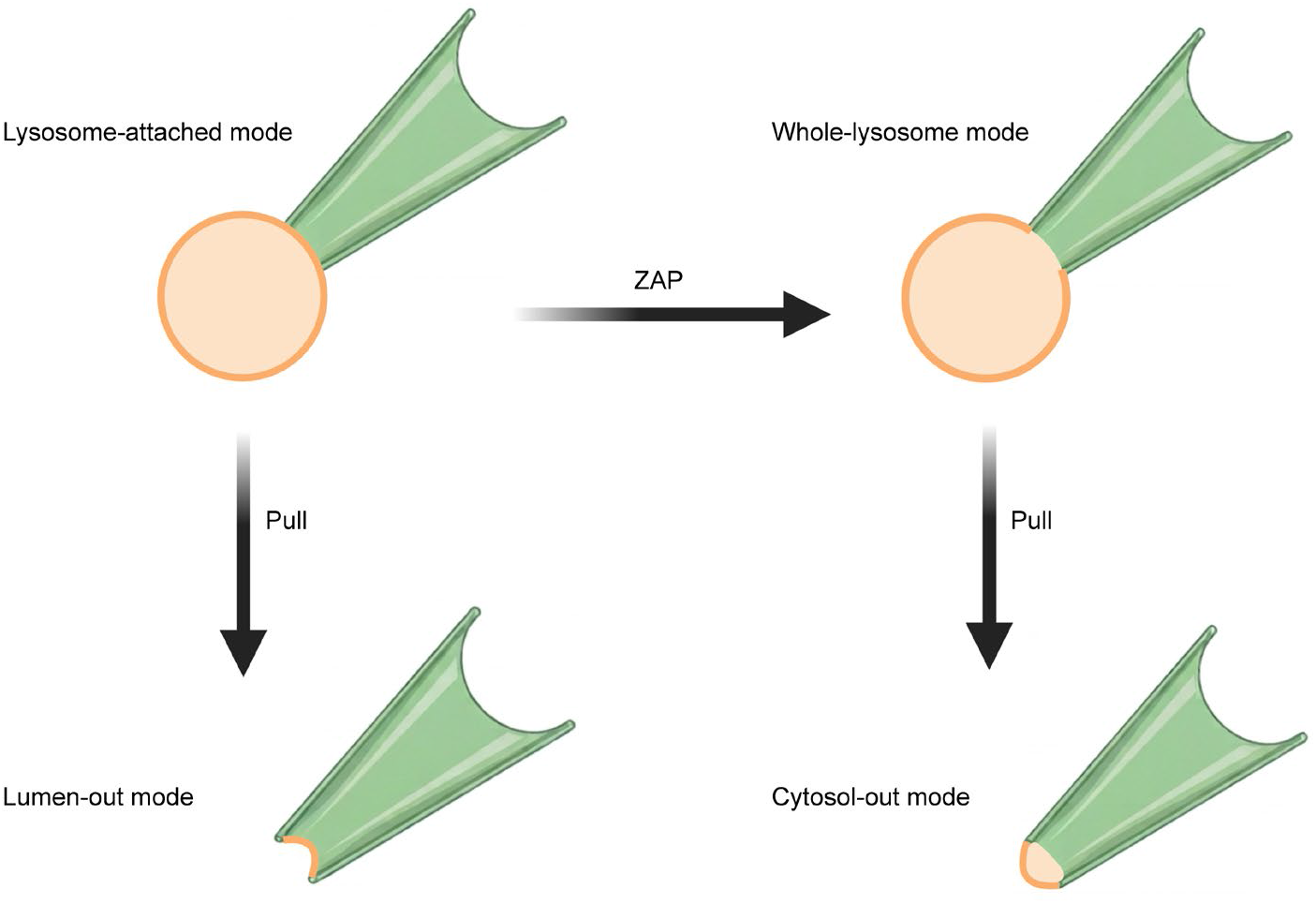
Single-lysosome recording configurations. Single-lysosome recordings can be performed in four configurations: endolysosome-attached, whole-lysosome, lumen-out, and cytosol-out modes. The endolysosome-attached and whole-endolysosome modes represent intact organelle recording configurations, whereas lumen-out and cytosol-out modes are obtained after membrane excision, exposing either the luminal or cytosolic side of the lysosomal membrane to the bath solution.

**NOTE:** In this configuration, the membrane potential of the endolysosome is unknown; therefore, the applied voltage does not reflect the absolute transmembrane potential.

**CRITICAL:** Current polarity is opposite to that observed in whole-endolysosome recordings.

B. **Lumen-out configuration (excised patch)** Quickly retract the pipette to excise the membrane patch from the vesicle.

**CRITICAL:** The excised patch may reseal into a vesicle; briefly remove the pipette from solution to rupture the outer membrane if necessary.

**NOTE:** This configuration exposes the luminal side of the membrane to the bath solution, allowing direct control of luminal conditions.

C. **Whole-endolysosome configuration** Apply a ZAP pulse (initially 25 µs; increase to 50 µs or longer duration if needed) to rupture the membrane patch and establish whole-endolysosome mode.

**CRITICAL:** Increase ZAP duration gradually (2-10x) before increasing voltage to avoid damaging the vesicle.

**NOTE:** Successful break-in is indicated by the appearance of capacitive transients and a decrease in series resistance.

D. **Cytosol-out configuration (excised patch from whole-endolysosome)** After establishing whole-endolysosome mode, gently retract the pipette to excise a membrane patch from the vesicle.

**CRITICAL:** Retraction should be rapid and controlled to obtain a stable excised patch.

**NOTE:** In this configuration, the cytosolic side of the membrane faces the bath solution, allowing direct control of cytosolic conditions.

### Recording protocols

#### Timing: 1-5 min per recording protocol (the duration varies)

After establishing the whole-endolysosome recording configuration, proceed with the corresponding recording protocol (**whole-endolysosome recordings**).

30. Compensate capacitance transients using amplifier circuitry until series resistance is <100 MΩ and capacitance is within 0.1-8 pF

**NOTE:** If capacitance compensation fails (e.g., no capacitance value is displayed or autocompensation error occurs), do not proceed with recording.

31. Select the recording mode (voltage-clamp recordings or current-clamp) and perform recordings as below.
  a. For voltage-clamp recordings:
    i. Set an appropriate holding potential depending on the channel and experimental conditions.
    ii. Apply voltage-step protocols using Clampex and record currents.

**NOTE:** Recordings are typically performed at room temperature unless otherwise specified.

b. For current-clamp recordings:
  i. Set I = 0 mode to disconnect external current input.

**CRITICAL:** Ensure the holding current is set to 0 before switching modes; non-zero holding current may disrupt the giga-seal.

ii. Load or design an appropriate current-clamp protocol.
iii. Switch to current-clamp (IC) mode.
iv. Monitor membrane potential and adjust tuning parameters to generate a stable sawtooth voltage response (~10 mV amplitude).
v. Enable automatic reduction of series resistance compensation if oscillations occur.
vi. Adjust pipette capacitance neutralization until slight overshoot appears.

**NOTE:** Overcompensation may induce oscillations and damage the vesicle; reduce slightly from the optimal value.

vii. Perform series resistance compensation (auto or manual adjustment).
viii. Acquire voltage recordings using the selected protocol.

For cytosol-out recordings (optional), proceed as follows: After completing the whole-endolysosome recording configuration, slowly retract the pipette to obtain a stable excised membrane patch to establish cytosol-out configuration (**single-channel recordings**).

**CRITICAL:** The membrane fragment should reseal at the pipette tip to form a patch with the cytosolic side exposed to the bath solution.

32. Set the headstage feedback resistor to 50 GΩ and enable headstage cooling to minimize noise.

**CRITICAL:** Turn on headstage cooling at least 1 h before recording to ensure signal stability.

33. Acquire single-channel currents using voltage-step protocols and verify channel activity.

**NOTE:** If no channel activity is observed, discard the recording and repeat with a new vesicle.

## Expected outcomes

Typical current traces are observed as the patch pipette enters the recording solution and approaches the target vesicle. Properly fabricated pipettes used for endolysosomal patch-clamp recordings typically exhibit resistances of 5-8 MΩ. Following seal formation, the seal resistance increases to the giga ohm range (typically 1-16 GΩ), indicating successful giga-seal formation.

Upon establishing the whole-endolysosome configuration, capacitive transients become apparent, reflecting electrical access to the vesicle lumen. From these transients, series resistance and membrane capacitance can be estimated. Under typical conditions, series resistance ranges from 10 to 100 MΩ, and capacitance ranges from 0.1 to 8 pF.

In whole-endolysosome recordings, membrane currents can be evoked using voltage-step protocols. These CLN3-mediated currents are typically in the picoampere (pA) range and are activated at positive voltages (Figure 4 A-C). Ramp Protocol (−150 to +150 mV): The current-voltage (I-V) relationship (Figure 4A) shows minimal current at negative potentials but a robust increase upon depolarization beyond 0 mV, demonstrating clear outward rectification. Step Protocol (−100 to +190 mV): Discrete current traces (Figure 4B-C) are observed only during positive voltage steps, confirming that the channel opens in response to membrane depolarization. High-quality recordings (achieved after a successful break-in at Step 29) must exhibit low baseline noise and minimal leak currents at hyperpolarized potentials. This confirms a stable Giga-seal and ensures the observed outward currents reflect true CLN3 activity.

**Figure 4.**
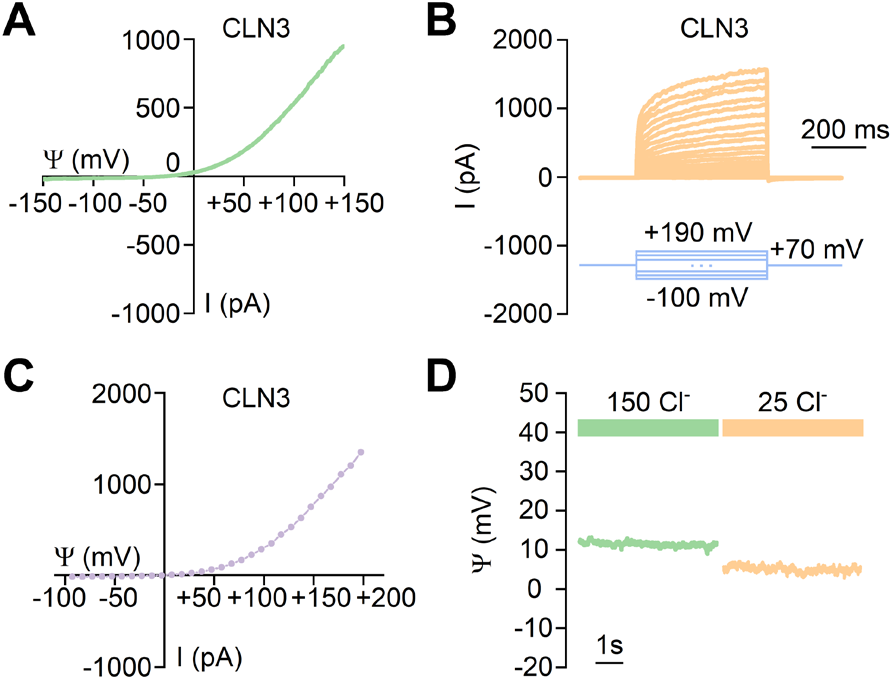
Representative whole lysosome recordings. (A) Representative current-voltage (I-V) relationship of CLN3-mediated currents, showing outward rectification over the tested voltage range. (B) Representative current traces recorded from CLN3 under step voltage protocols (−100 mV to +190 mV), illustrating time-dependent current activation. Scale bar, 200 ms. (C) Averaged I-V relationship of CLN3 currents compiled from multiple recordings, confirming the voltage-dependent current profile. (D) Effect of luminal chloride concentration on CLN3 activity. Representative current traces recorded under symmetric conditions of 150 mM Cl^−^ (green) and reduced luminal chloride (25 mM Cl^−^, orange), showing altered current amplitude.

In current-clamp mode, changes in membrane potential can be monitored in response to alterations in ionic compositions (Figure 4D). For instance, shifting the cytosolic-side chloride concentration from 150 mM to 25 mM induces a dynamic change in membrane potential. In excised or attached patch configurations, single-channel activity may be observed as discrete step-like current transitions between closed and open states. These events are characterized by defined amplitude levels and reproducible kinetics, provided that the signal-to-noise ratio is sufficiently high.

## Limitations

This protocol requires pharmacological or genetic enlargement of endolysosomes, which may alter membrane composition, lipid distribution, or regulatory protein interactions. Vacuolin-1 and YM201636 may affect trafficking pathways or phosphoinositide balance, potentially influencing channel behavior. Mechanical isolation introduces selection bias toward larger and more mechanically stable vesicles. Smaller or more fragile endolysosomes may be underrepresented in recordings. Whole-endolysosome configuration allows washout of luminal or cytosolic modulators, which may affect channel gating over time. Regulatory proteins or second messengers associated with lysosomal membranes may be diluted during recording. This protocol is technically demanding and requires extensive optimization of pipette fabrication and micromanipulation. Very small organelles or complex membrane systems such as the endoplasmic reticulum cannot be reliably accessed using this approach.

## Troubleshooting

### Problem 1

Enlarged lysosomes are not observed after Vacuolin-1, YM201636 or Wortmannin + Latrunculin B treatment (related to Step 6).

### Potential solution

Verify drug concentration and incubation time. Optimize concentration (between 0.5-5 μM) depending on cell type. Confirm cell viability and ensure cells are 40-70% confluent at time of treatment. Test an alternative enlargement strategy (e.g., Rab5Q79L expression).

### Problem 2

Isolated vesicles rupture during mechanical dissection (related to Steps 16-20).

### Potential solution

Reduce mechanical force applied during extrusion. Select vesicles positioned near the cell periphery. Use a smoother and slightly blunter isolation pipette tip. Ensure cells are firmly attached to poly-L-lysine coated coverslips.

### Problem 3

Giga-seal cannot be established (related to Steps 21-29).

### Potential solution

Confirm that pipette tips are properly fire-polished and not excessively sharp. Reduce the positive pressure prior to contact with the cell. Ensure that the bath solution is clean and free of debris (e.g., by filtration). Replace the pipette if its resistance falls outside the range of 5-8 MΩ.

### Problem 4

Break-in fails after ZAP pulse (related to Step 29).

### Potential solution

Increase ZAP duration incrementally (50-100 μs). Avoid excessive positive pressure prior to break-in. Ensure vesicle diameter is ≥3 μm.

### Problem 5

Seal becomes unstable during recording (related to Steps 30-31).

### Potential solution

Move vesicle slightly upward after giga-seal formation to stabilize geometry. Minimize bath turbulence. Avoid excessive solution exchange during recording.

## Resource availability

### Lead contact

Further information and requests for resources and reagents should be directed to and will be fulfilled by the lead contact, Dr. Lily Yeh Jan.

### Technical contact

Technical questions on executing this protocol should be directed to and will be answered by the technical contact, Dr. Yayu Wang.

### Materials availability

This study didn’t generate new unique reagents.

### Data and code availability

This study did not generate or analyze datasets/code.

## Acknowledgments

We thank all members of the Jan lab for valuable discussions. This work was supported by the NIH grant to L.Y.J. (R35NS122110).

## Author contributions

Y.W. performed both the experiments and analyses. L.Y.J. and Y.W. wrote and edited the manuscript.

